# Developmental atlas of white lupin cluster roots

**DOI:** 10.1101/2020.03.26.009910

**Authors:** Cécilia Gallardo, Bárbara Hufnagel, Alexandre Soriano, Fanchon Divol, Laurence Marquès, Patrick Doumas, Benjamin Péret

## Abstract

During the course of evolution, plants have developed various strategies to improve micronutrient acquisition, such as cluster roots. These spectacular structures are dedicated to efficient phosphate remobilization and acquisition. When exposed to Pi-limitation, white lupin forms cluster roots made of dense clusters of short specialized roots, called rootlets. Although the physiological activity of rootlets has been well studied, their development remains poorly described. Here, we provide a developmental atlas of white lupin early rootlet development, using molecular markers derived from the model plant Arabidopsis. We first focused on cell division patterns to determine which cells contribute to the rootlet primordium. Then, we identified homologs of previously described tissue specific genes based on protein sequence analysis and also using detailed transcriptomic data covering rootlet development. This study provides a comprehensive description of the developmental phases of rootlet formation, highlighting that rootlet primordium arises from divisions in pericycle, endodermis and cortex. We describe that rootlet primordium patterning follows eight stages during which tissue differentiation is established progressively.

**Highlight:** White lupin cluster roots consist in the formation of numerous rootlets whose development can be divided in 8 stages and involves divisions in the pericycle, endodermis and cortex.

## Introduction

Cluster roots (CRs) are considered to be one of the major adaptations to improve plant nutrient acquisition, together with nitrogen-fixing nodules and mycorrhizae (Neumann and Martinoia, 2002). However, CRs differ from nodules and mycorrhizae as they form without the presence of a symbiotic partner (Lamont, 2003). CR bearing species can be found in soils where nutrients are poorly available (Dinkelaker *et al*., 1995; Lambers *et al*., 2003). Plants forming CRs can absorb inorganic phosphate (Pi) at a faster rate than non-forming CRs plants and access to a larger pool of Pi due to increased soil exploration (Vorster and Jooste, 1986). Because this developmental adaptation has a selective advantage, CR arose in a whole range of distant families from Cyperaceae and Restionaceae in monocots to Betulaceae, Casuarinaceae, Cucurbitaceae, Eleagnaceae, Fabaceae, Moraceae, Myricaceae and Proteaceae in dicots (Dinkelaker *et al*., 1995; Lambers *et al*., 2003). Interestingly, most of these families have lost the ability to form mycorrhizae (Oba *et al*., 2001; Delaux *et al*., 2014; Maherali *et al*., 2016).

CR is an adaptive trait of plants to cope with P-depleted soils (Neumann and Martinoia, 2002; Lambers *et al*., 2015). Because phosphate is mostly concentrated in the topsoil layer, CRs are mainly produced in the upper part of the root system (Lynch and Brown, 2001). Hereafter, CRs will refer to the entire length of the lateral root that comprises at least one cluster of tightly grouped third-order roots termed rootlets (Dinkelaker *et al*., 1995; Skene, 1998). CRs are an exacerbated developmental response that results from massive induction of rootlets and represent a good model to study root adaptive developmental responses to abiotic constraints. White lupin (*Lupinus albus*) is one of only few crops that can form CRs (Fig. 1A-C). It has received most attention for studies on CR morphology and physiology (Johnson *et al*., 1996; Watt and Evans, 1999; Hagström *et al*., 2001; Neumann and Martinoia, 2002; Uhde-Stone *et al*., 2003; Cheng *et al*., 2011). White lupin (WL) can form up to 35-45 rootlets per cm (Dinkelaker *et al*., 1989) and secrete massive amount of organic acids and protons (Massonneau *et al*., 2001; Sas *et al*., 2001; Neumann and Martinoia, 2002). Such physiological modifications can increase the Pi availability of soil, resulting in a beneficial effect to other species in mixed cultures (Braum and Helmke, 1995; Cu *et al*., 2005; Li *et al*., 2010). For instance, mixed culture of wheat and white lupin was shown to increase shoot Pi uptake up to 45 % in wheat (Cu *et al*., 2005). This shows how white lupin can be used to improve the poor efficiency of Pi uptake by several crops for a sustainable agriculture perspective.

**Fig. 1.**
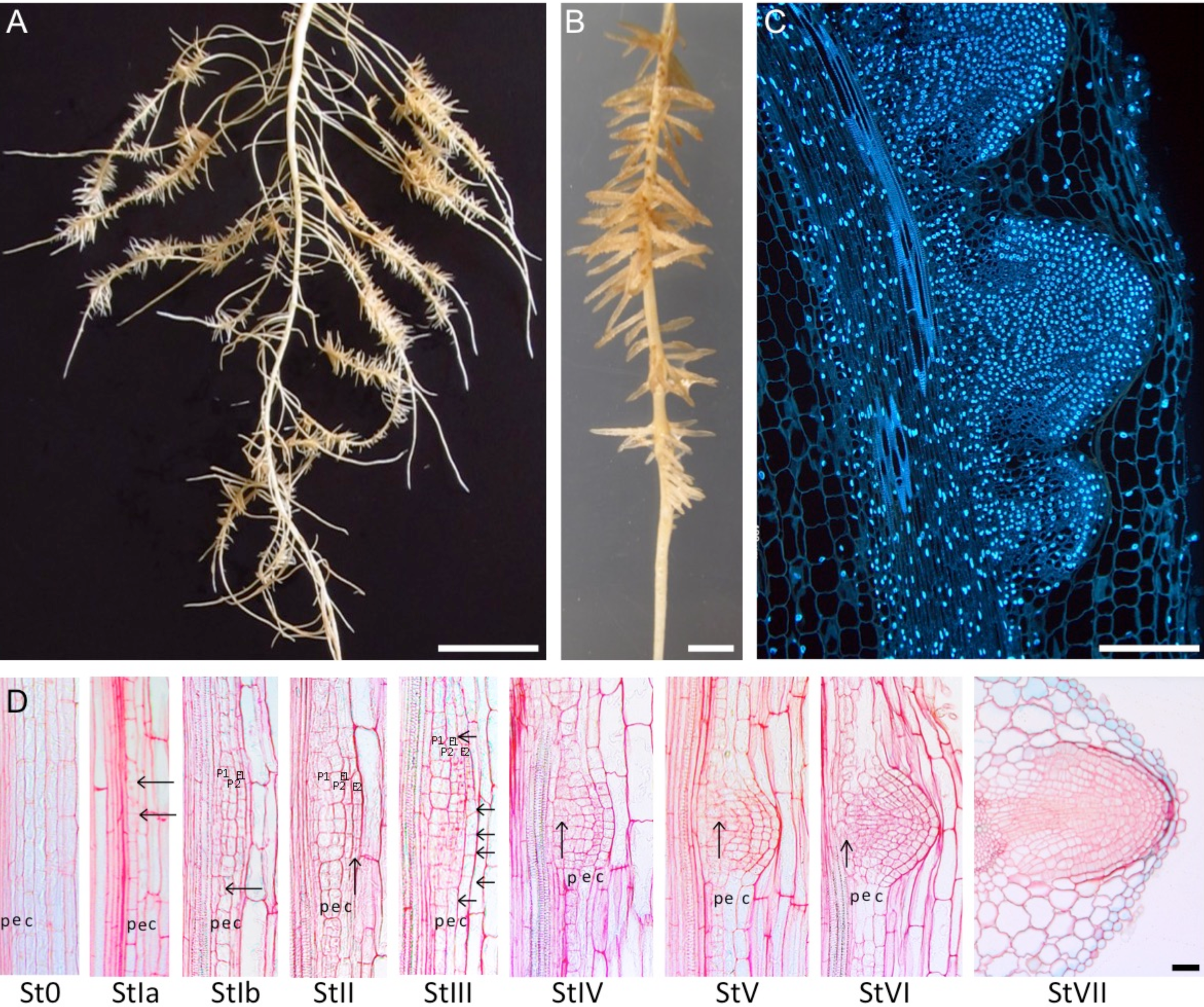
Histological description of the 8 stages of rootlet primordium formation. (A) Whole root system of a 21-day old white lupin in phosphate deficiency conditions harbouring numerous cluster roots. (B) One cluster root harbouring numerous rootlets. (C) Nuclear staining (DAPI) of rootlet primordium during the emergence stage. (D) Longitudinal sections of a cluster root depicting all developmental stages of rootlet development (ruthenium red stained). Last image (StVII) is a cross section. St0: Stage 0. Parental tissues before visible events. StIa: Stage Ia. First asymmetric division in the pericycle. StIb: Stage Ib. First periclinal division in the pericycle and first anticlinal division in the endodermis. StII: Stage II. Formation of a flattened four-layered primordium: two pericycle layers and two endodermal layers. StIII: Stage III. Several rounds of extra anticlinal divisions in the endodermis and first cortical divisions. StIV: Stage IV. Divisions rate increase and start to shape a dome. StV: Stage V. The dome shape is now well established and the rootlet primordium passes through the cortical layers. StVI: Stage VI. The primordium is now half-way through the cortical layers. StVII: Stage VII. The rootlet primordium is about to emerge from the maternal root. p: pericycle, e: endodermis, c: cortex. Arrows pinpoint key division planes. Scale bars: 1 cm (A), 1mm (B), 200 µm (C), 50 µm (D).

In many angiosperms, secondary roots (or lateral roots) arise from pericycle as described in the model plant *Arabidopsis thaliana* (Laskowski *et al*., 1995; Malamy and Benfey, 1997; Orman-Ligeza *et al*., 2013; Herrbach *et al*., 2014). In some monocots, lateral roots show contribution of dividing endodermal cells including some Poaceae like rice (*Oriza sativa*) and maize (*Zea mays*) (Hochholdinger *et al*., 2004). In dicots, some Legume species also show contribution of division in endodermis and cortex including soybean (*Glycine max*), *Lotus japonicus* and alfalfa (*Medicago truncatula*) (Byrne *et al*., 1977; Op den Camp *et al*., 2011; Herrbach *et al*., 2014). In Arabidopsis, it was proposed that lateral root primordium development proceeds through an 8-stage process (Malamy and Benfey, 1997) that can be summed up as a two-step developmental model (Laskowski *et al*., 1995). Early developmental phase of lateral root primordium (stage I-IV) starts with the first division in two adjacent pericycle cells followed by several rounds of divisions to produce a four-layered primordium (Malamy and Benfey, 1997). Then, cells start to differentiate and acquire various identities as tissues organize around a meristematic zone (stage V-VII) (Trinh *et al*., 2018). Transition between these two developmental phases marks the onset of quiescent centre establishment and formation of meristematic initials (Goh *et al*., 2016). With regards to these processes, very little information is available for third-order roots, both in Arabidopsis and lupin. In WL, it was described that xylem pole pericycle cells are recruited to become founder cells (Johnson *et al*., 1996) and a predominant role for auxin was demonstrated (Gallardo *et al*., 2019). However, a full description of the entire developmental process of rootlet formation is still missing.

In an effort to generate a developmental atlas of rootlet formation in white lupin, we performed an anatomic study to define discrete developmental stages throughout early rootlet development. Since the root structure of WL is more complex than Arabidopsis root, it appeared important to use a set of various molecular markers to identify division and differentiation events. We first used the cell division marker *CYCLIN B1;1* (*CYCB1;1*) from Arabidopsis whose promoter revealed to drive expression in WL dividing cells. This allowed us to identify cell division in pericycle but also in endodermal and cortical cells during rootlet formation. In a second approach, we decided to take advantage of a set of tissular markers and identify their homolog genes in WL genome. We combined sequence analysis to detailed expression profiling to select the best candidates amongst WL highly duplicated gene families. We generated transgenic hairy root plants expressing these tissular markers that show a strong conservation of tissular specificity. Using this approach, we provide a developmental atlas of cell division and differentiation during white lupin rootlet primordium development.

## Materials and methods

### Plant materials and growth conditions

Seeds of white lupin (*Lupinus albus L. cv. Amiga*) calibrated at 7 mm were used in all experiments. White lupin plants were cultivated in growth chambers under controlled conditions (16 h light / 8 h dark, 25°C day / 20°C night, 65 % relative humidity, and PAR intensity 200 μmol.m^-2^.s^-1^) in hydroponic conditions. The hydroponic solution was modified from (Abdolzadeh *et al*., 2010) without phosphate, and was composed of: MgSO_4_ 54 μM; Ca(NO_3_)_2_ 400 μM; K_2_SO_4_ 200 μM; Na-Fe-EDTA 10 μM; H_3_BO_3_ 2.4 μM; MnSO_4_ 0.24 μM; ZnSO_4_ 0.1 μM; CuSO_4_ 0.018 μM; Na_2_MoO_4_ 0.03 μM. Plants were grown either in 1,6 L pots or 200 L tanks. Solution was aerated continuously. For plants in pots, the nutrient solution was renewed every 7 days.

### Molecular cloning

The primers to amplify the promoter sequences of *LaSCR1, LaWOL, LaPEP, LaEXP7* were designed using Primer3plus (http://www.bioinformatics.nl/cgi-bin/primer3plus/primer3plus.cgi). All primer sequences used are summarized in Table S1. Primers were used to amplify at the minimum 1500 bp upstream of the start codon from white lupin genomic DNA. Sizes of all the promoters amplified are summarized in Table S1. AttB1 (5’-GGGGCCAAGTTTGTACAAAAAAGCAGGCT-3’) and attb2 (5’-CCCCCCACTTTGTACAAGAAAGCTGGGT-3’) adapters were added to the primers. Amplified fragments were subsequently cloned into the pDONR221 by Gateway reaction. The promoters were then cloned into the binary plasmid pKGWFS7 containing a GFP-GUS fusion by Gateway cloning.

### Bacterial strain

*Agrobacterium rhizogenes* strain *ARquaI* was used to perform *hairy root* transformation of white lupin. Bacteria were transformed with the binary plasmid by electroporation. Transformation was confirmed by PCR and sequencing. LB agar plates added with sucrose 2%, acetosyringone 100 μM and appropriate antibiotics were inoculated with 200 μL of liquid bacteria culture, and incubated at 28°C for 24 h to get a bacterial lawn. Bacterial lawn was used for white lupin seedling transformation.

### Hairy root transformation of white lupin

White lupin hairy root transformation was performed as previously described (Uhde-Stone *et al*., 2005). White lupin seeds caliber 8 mm were surface sterilized by 4 washes in osmosis water, 30 min sterilization in bleach (Halonet 20%) and washed 6 times in sterile water under sterile conditions. Seed were germinated on half MS medium (pH was adjusted to 5.7). After germination, radicles of 1 cm were cut over 0.5 cm with a sterile scalpel. The radicles were inoculated with the *Agrobacterium rhizogenes* lawn. Fifteen inoculated seedlings were placed on square agar plates (0.7 % agar in 1X Hoagland solution) containing 15 μg.mL^-1^ Kanamycin. Composition of Hoagland medium without phosphate was the following one: MgSO_4_ 200 μM; Ca(NO_3_)_2_ 400 μM; KNO_3_ 325 μM; NH_4_Cl 100 μM; Na-Fe-EDTA 10 μM; H_3_BO_3_ 9.3 μM; MnCl_2_ 1.8 μM; ZnSO_4_ 0.17 μM; CuSO_4_ 0.06 μM; Na_2_MoO_4_ 2.3 μM. Plates were placed vertically in controlled conditions: 16 h light / 8 h dark, 25°C day / 20°C night, 65 % relative humidity, and PAR intensity 200 μmol.m^-2^.s^-1^. After 7 days on plates, 60 seedlings were transferred to 12×16.5×5.5 cm trays (20 seedlings per tray) and watered with 500 mL pure water. After 12 days, plants presenting hairy roots were transferred to hydroponics in 1.6 L pots containing nutrient solution. Nutrient medium was renewed each week. After 7 days in hydroponic conditions, CRs were sampled on *hairy root* plants.

### Histochemical analysis

Histochemical staining of β-glucuronidase was performed on CRs from *hairy root* plants. Samples were incubated in a phosphate buffer containing 1 mg.mL^-1^ X-gluc as a substrate (X-Gluc 0.1 %; phosphate buffer 50 mM, pH 7, potassium ferricyanide 2 mM, potassium ferrocyanide 2 mM, Triton X-100 0.05 %). Zone of CRs with non-emerged rootlets were cutted into 4 to 5 sections and fixed (formaldehyde 2%, glutaraldehyde 1%, caffeine solution 1%, phosphate buffer at pH 7). Fixation was performed for 2.5 h under agitation at room temperature and then 1.5 h at 4°C.

### Microscopy analysis

To generate thin sections, roots were progressively dehydrated in ethanol solutions with increased concentrations: 50% (30 min), 70% (30 min), 90% (1 h), 95% (1 h), 100% (1 h), 100% (overnight). Samples were impregnated with 50% pure ethanol and 50% resin (v/v) for 2 days, then in 100 % resin for 5 days. CRs were embedded in Technovit 7100 resin (Heraeus Kulze, Wehrheim, Germany) according to the manufacturer’s recommendations. Thin sections of 10 μm were produced using a microtome (RM2165, Leica Microsystems) and counterstained for 30 min with 0.1% ruthenium red and rinsed two times with ultrapure water. For nuclei visualization, 10 μm thick cluster roots sections were stained for 10 min with 3 μM DAPI solution in PBS buffer at room temperature. After washing root three times with PBS buffer for 5 min, the section were mounted in PBS buffer and analyzed. An argon laser at 405 nm provided excitation for DAPI staining. The fluorescence emission signal was detected using a band-pass filter of 420-480 nm for DAPI. Imaging was performed in Montpellier RIO imaging Platform (http://www.mri.cnrs.fr/fr/) with a stereomicroscope (SZX16, Olympus) for macroscopic root images, an epifluorescence microscope with a colour camera (Olympus BX61 with Camera ProgRes^®^C5 Jenoptik and controlled by ProgRes Capture software) for thin root section, and a confocal microscope (Leica SP8, Leica Microsystems, https://www.leica-microsystems.com/fr/) for nuclei visualization.

### Phylogenetic trees

White lupin cDNA-deduced protein sequences were retrieved from the available white lupin genome sequence tool (https://www.whitelupin.fr/). Arabidopsis protein sequences were extracted from TAIR database (https://www.arabidopsis.org) and protein sequences from *Lupinus angustifolius, Medicago truncatula, Cicer arietinum, Glycine max* were downloaded from NCBI database (https://www.ncbi.nlm.nih.gov/guide/proteins/). Phylogenetic trees were constructed using the phylogeny tool NGPhylogeny.fr (https://ngphylogeny.fr/) (Lemoine *et al*., 2019). The bootstrap consensus tree was inferred from 500 replicates. Branches corresponding to partitions reproduced in less than 50% bootstrap replicates were collapsed. Sequences were aligned using MUSCLE 3.7 and Gblocks with default parameters were used to eliminate poorly aligned positions and divergent regions. PhyML 3.0 was used to perform phylogenetic inference, using SH-like as aLRT test. Tree visualization was performed through the iTOL v4.4.2 platform (Letunic and Bork, 2019).

## Results

### Describing the developmental stages of rootlet primordium

To provide a detailed description of cluster root development, thin longitudinal sections of 7 to 9 day-old lupin seedlings were stained with ruthenium red to observe all developmental stages. Early events of rootlet development could be described in 8 discrete developmental stages (Fig. 1D).

#### Stage I

Initiation is the first visible event of CR formation. It consists of two asymmetrical anticlinal divisions in the pericycle (Fig. 1D StIa). Initiation continues with the appearance of divisions in parallel orientation compared to the root axis (Fig. 1D StIb, black arrow). In the longitudinal plane, close to 6 cells show these periclinal divisions. Peripheral cells are not dividing, creating the boundaries of the primordium. In the overlaying endodermis, an increased number of anticlinal divisions is clearly seen as compared to the surrounding endodermal cells. About 8 cells are formed from these divisions.

#### Stage II

All pericycle cells participating in the rootlet primordium seem to have divided. These divisions in the pericycle lead to the formation of two layers, named P1 (inner layer) and P2 (outer layer). Following these divisions, cells start to swell and expand in the radial direction. Simultaneously, periclinal divisions occur in the endodermis that divides this layer into E1 (inner layer) and E2 (outer layer) (Fig. 1D StII, black arrow).

#### Stage III

A dozen of cells continue to divide in the 4-layered primordium. Anticlinal divisions are seen in the most peripheral cells in the endodermis. At the same time, anticlinal divisions are also seen in the cortex (Fig. 1D StIII, purple arrows).

#### Stage IV

A typical dome shaped rootlet primordium begins to form. The dividing cortex comprises 10 cells in length. Four cells at the base of the primordium started to divide between the xylem pole and the P1 inner layer (Fig. 1D StIV, black arrow) giving rise to the future vasculature. Numerous radial, anticlinal and periclinal divisions happen in the tissues derived from pericycle, endodermis and cortex.

#### Stage V

Primordium expands radially, pushing the overlaying cortical layers. Cells at the base of the primordium (Fig. 1D StV, black arrow) continue to divide. The number of cells in the overlaying cortex increases to 11 cells in length and are now forming part of the tip of the primordium.

#### Stage VI

The rootlet primordium is now strongly deforming the cortical layers around. The stage VI primordium begins to look like a mature root with several layers, although a typical meristematic organization is not clear yet due to the apparent absence of a quiescent centre. In the absence of tissue-specific markers, it is difficult to make strong assumption regarding the nature of each tissue. The presence of elongated cells at the base of the primordium seems to indicate the onset of the vasculature.

#### Stage VII

The rootlet primordium is about to cross the last layer of the cluster root, the epidermis. As the primordium enlarges, distinguishing the different tissues inside the growing rootlet becomes challenging due to the high number of cells. Many anticlinal divisions seem to continue in the different layers of the primordium. The vascular tissue seems to be now fully established as many elongated cells are connecting the cluster root stele to the rootlet primordium.

#### Stage VIII

No visible difference can be seen in the cellular organization of the primordium but cell elongation pushes the primordium beyond the epidermis and the primordium reaches the rhizosphere.

### Following cell divisions during rootlet primordium formation

Although many cell divisions can be seen from cell shape and the appearance of extra division planes, the use of a molecular division marker allows to easily monitor mitotic activity and define dividing cells at an early stage. The promoter of the *Arabidopsis thaliana CYCB1;1* gene is activated during the G2/M transition (Colón-Carmona *et al*., 1999). We transformed WL roots with a construct containing the Arabidopsis *CYCB1;1* promoter fused to the ß-glucuronidase reporter gene (pAt*CYCB1;1:GUS*). This marker shows expression in the cluster root apex, very similar to the expression described in Arabidopsis, suggesting a conservation of the expression during cell cycle (Supplementary Figure S1A). Similarly, expression is visible at early stages of rootlet initiation and throughout the developmental process up to their emergence (Supplementary Figure S1B-D). We next produced thin cross sections to analyse in detail the mitotic activity in cells of the rootlet primordia. We observed that the first visible event of rootlet initiation is the activation of the pAt*CYCB1;1:GUS* marker in one pericycle cell in front of each xylem pole (Fig. 2A), before division is visible from an anatomical point of view, defining Stage 0. Subsequently, one or more divisions are seen in the pericycle cells (Fig. 2B,C) suggesting that more than one cell participate in the creation of the primordium. Then, divisions are also seen in the endodermis (Fig. 2C) and cortex (Fig. 2D). These divisions seem mainly periclinal and therefore produce extra layers of these tissues. Once the rootlet primordium reaches emergence, several divisions can be seen in what seems to be a fully dividing meristematic zone (Fig. 2E).

**Fig. 2.**
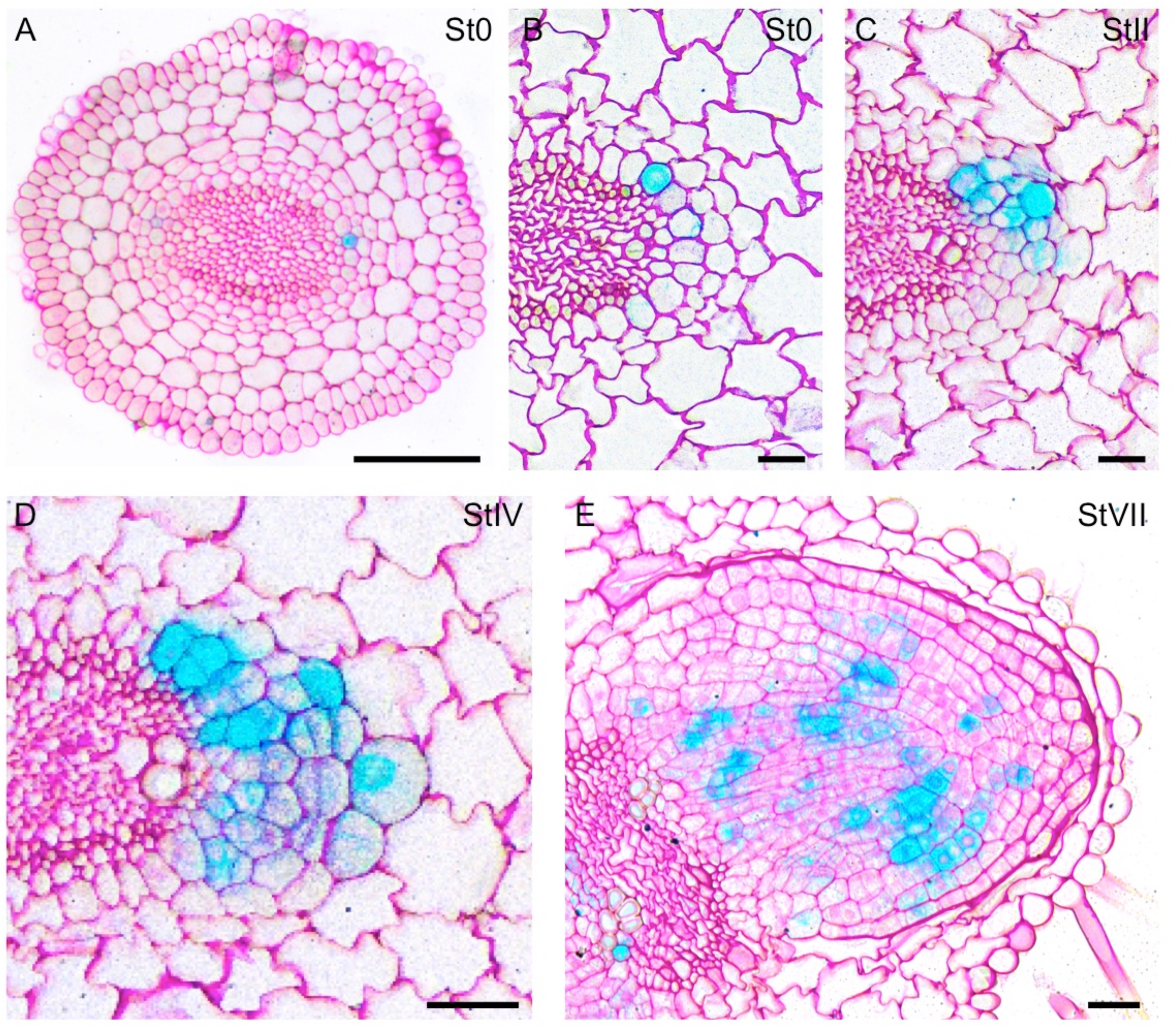
Cell divisions during rootlet formation marked by the Arabidopsis *pAtCYCB1;1:GUS* marker. (A-E) Thin cross sections of hairy root transformed cluster roots expressing the *pAtCYCB1;1:GUS* marker of the G2/M transition of the cell cycle. (A) Stage 0: one pericycle cell in front of each xylem pole expresses the marker. (B) Close-up view of the earliest observable event of rootlet initiation depicting one cell preparing to divide. (C) Stage II: active divisions in the pericycle and endodermis. (D) Stage IV: numerous divisions occur in the pericycle, endodermis and cortex. (E) Stage VII: just prior to emergence, numerous divisions are seen in what appears to be a fully functional and organized meristem. Scale bars are 100 µm (A) and 50 µm (B-E).

### Expression of tissue-specific markers in CR

Classic histology allows to describe precisely the early developmental stages of rootlet formation and to define discrete stages associated with particular anatomic features. However, cell types cannot be easily identified based solely on their shape during early developmental stages. To which extent pericycle, endodermis and cortex tissues are involved in rootlet initiation and rootlet outgrowth has yet to be determined. Another matter is to determine when tissues are starting to differentiate. To address these questions, a limited number of molecular markers (using the ß-glucuronidase gene as reporter) specifically expressed in different tissue types were tested. The corresponding white lupin genes were chosen (1) based on the protein sequence homology between known molecular markers in the model plant *Arabidopsis thaliana* and (2) based on their expression profile in the white lupin transcriptomic dataset. Due to genome triplication, gene families are often larger in white lupin than in Arabidopsis (Hufnagel *et al*., 2020) and expression data helped directing our choice.

First, we chose a list of molecular markers based on their tissular expression in *Arabidopsis thaliana*: two endodermal markers (WOL, SCR1), one cortical markers (PEP) and one epidermal marker (EXP7) (Sbabou *et al*., 2010; Marquès-Bueno *et al*., 2016; Sevin-Pujol *et al*., 2017). Orthologous lupin genes were found by comparing Arabidopsis protein sequences with all the cDNA-deduced protein sequences of WL genome. Comparison was made by multiple comparison alignment by local BLAST on the white lupin database and by generating phylogenetic trees containing the Arabidopsis sequence as well as Legume relatives: *Medicago truncatula, Cicer arietinum, Lupinus angustifolius* and *Glycine max* (Fig. 3). On the other hand, expression profiles of the selected WL genes were checked in a transcriptomic RNAseq database describing the spatial portions of a 5 days-old cluster root and therefore covering the different developmental stages from S0 to S7 (Fig. 4A) (Hufnagel *et al*., 2020). All genes were found to be expressed during rootlet development and genes were discriminated on their level of expression. For each gene family (3 to 4 genes), one gene was found to be more highly expressed that the others and was selected assuming that high gene expression would translate into a nicely visible marker (Fig. 4B-D). As a result, we selected the following WL genes for further analysis: *Lalb_Chr04g0258751* (WOL), *Lalb_Chr19g0123861* (SCR1), *Lalb_Chr11g0065071* (PEP) and *Lalb_Chr09g0324651* (EXP7). Gene sequence and expression data can be found on the White Lupin genome portal (https://www.whitelupin.fr/). The WL promoters (at least 1500 bp upstream start codon) of these genes were cloned upstream the ß-glucuronidase (GUS) sequence and introduced into WL roots via *Agrobacterium rhizogenes*-mediated root transformation (Uhde-Stone *et al*., 2005).

**Fig. 3.**
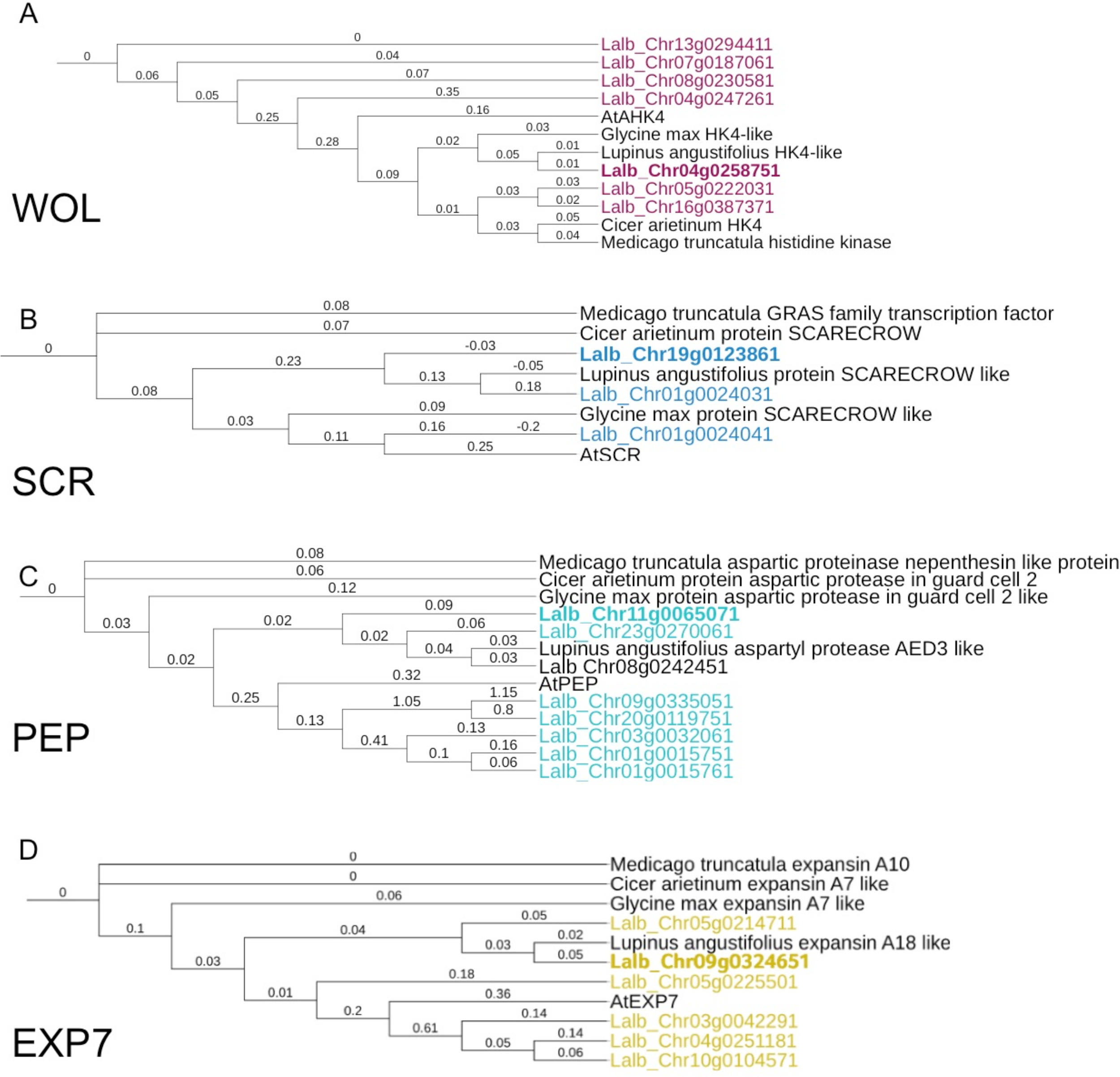
Maximum likelihood phylogenetic tree of white lupin orthologs and Arabidopsis gene markers. The trees were constructed using the phylogeny tool from NGPhylogeny.fr (https://ngphylogeny.fr/). Branches lengths are displayed. The orthologs selected to further experiments are highlighted. Analyses were performed using protein sequences extracted from white lupin genome (https://www.whitelupin.fr/) and from public database TAIR (https://www.arabidopsis.org) belonging to the model plant *Arabidopsis thaliana* and to the following Legume species: *Medicago truncatula, Cicer arietinum, Lupinus angustifolius* and *Glycine max*.

**Fig. 4.**
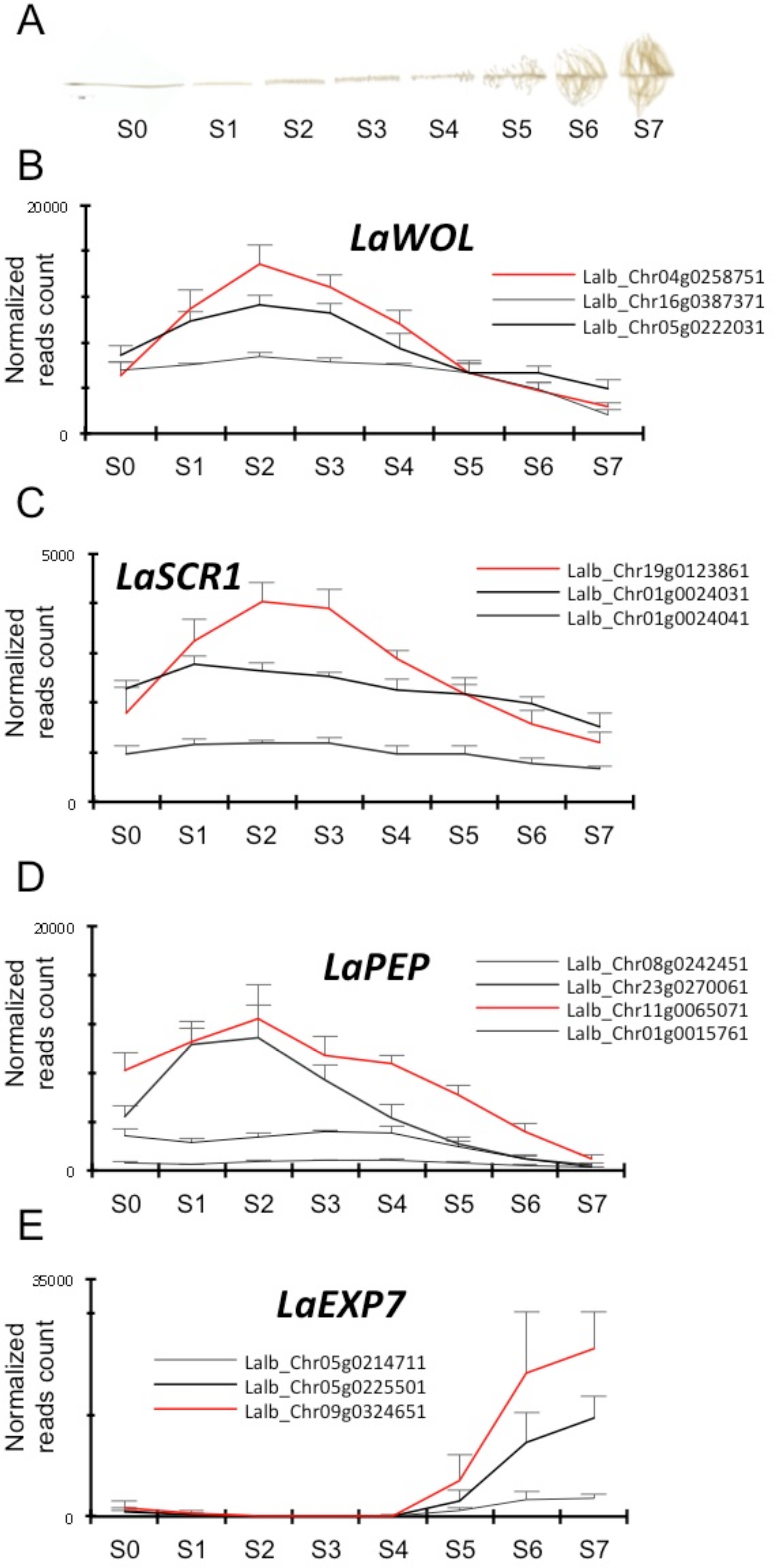
Candidate marker gene expression across rootlet development. (A) Data coming from transcriptomic temporal dataset of 8 developmental stages of rootlet formation as described in (Hufnagel *et al*., 2020). (B-E) Expression profile of white lupin ortholog genes to Arabidopsis tissue specific markers. Red lines indicate genes for which promoters were cloned for further studies. Data are mean ± SD of four biological replicates coming each from ten cluster roots sampled across 5 lupin plants (n=10).

To characterize tissue specific expression, their expression pattern was first assessed into secondary-order roots *i.e.* apex of cluster roots. CRs stained for GUS for each marker are shown in Fig. 5, as well as cross sections produced in these lines. The *pLaWOL:GUS* line was expressed in the stele of CR (Fig. 5A-B). Staining was not observed in the elongation zone but was seen in the differentiation zone in the stele (Fig. S2A). A transverse section through the differentiation zone shows GUS staining in stellar tissue along xylem axis and pericycle cells facing the xylem poles (Fig. 5B). Staining in the stele disappears in the primordium patterning zone and became restricted only to pericycle cells and the stele of the primordium. (Fig. S2B). *pLaWOL:GUS* driven expression was observed in the stele of emerged rootlet with a stronger staining in the meristematic zone were QC is expected to locate (Fig. S2C-E). Expression of *pLaSCR1:GUS* in cluster roots matched previous reports (Sbabou *et al*., 2010) and was strongly specific of the endodermis of CR tip (Fig. 5C,D). Promoter expression was also observed early on in rootlet primordia (Fig. S2F,G) and more specifically in the rootlet meristematic zone after their emergence (Fig. S2H-I). Expression of the *pLaPEP:GUS* construct was specific of cortical cells (Fig. 5E,F). Promoter is strongly expressed in the root meristematic zone, and is not expressed in endodermis/cortex initial and the QC (Fig. 5E). A transverse section in the beginning of elongation zone is shown in Fig. 5F showing the specificity of the cortical expression. Expression tends to fade away in the elongation zone in the shootward direction (not shown). Similar pattern for *pLaPEP:GUS* was observed in primordium and emerged rootlet (Fig. S2K-O). Expression of the *pLaEXP7:GUS* construct was absent from the very tip of the CR meristem and started at the onset of the elongation zone (Fig. 5G). A cross section performed in the differentiation zone showed the specificity of the epidermal expression (Fig. 5H). Staining was still present in the epidermal cells in the primordium patterning zone (Fig. S2P) but was absent from primordium (Fig. S2Q) and young rootlets (Fig. S2R,S). Expression resumes in older rootlets when elongation zone is fully active (Fig. S2T). The study of the 4 selected marker genes suggested that 3 of them could be used to further analysis during early rootlet primordium development (since *LaEXP7* shows no expression). We therefore selected *pLaWOL:GUS, pLaSCR1:GUS* and *pLaPEP:GUS* to provide a detailed description of rootlet developmental stages and tissue differentiation.

**Fig. 5.**
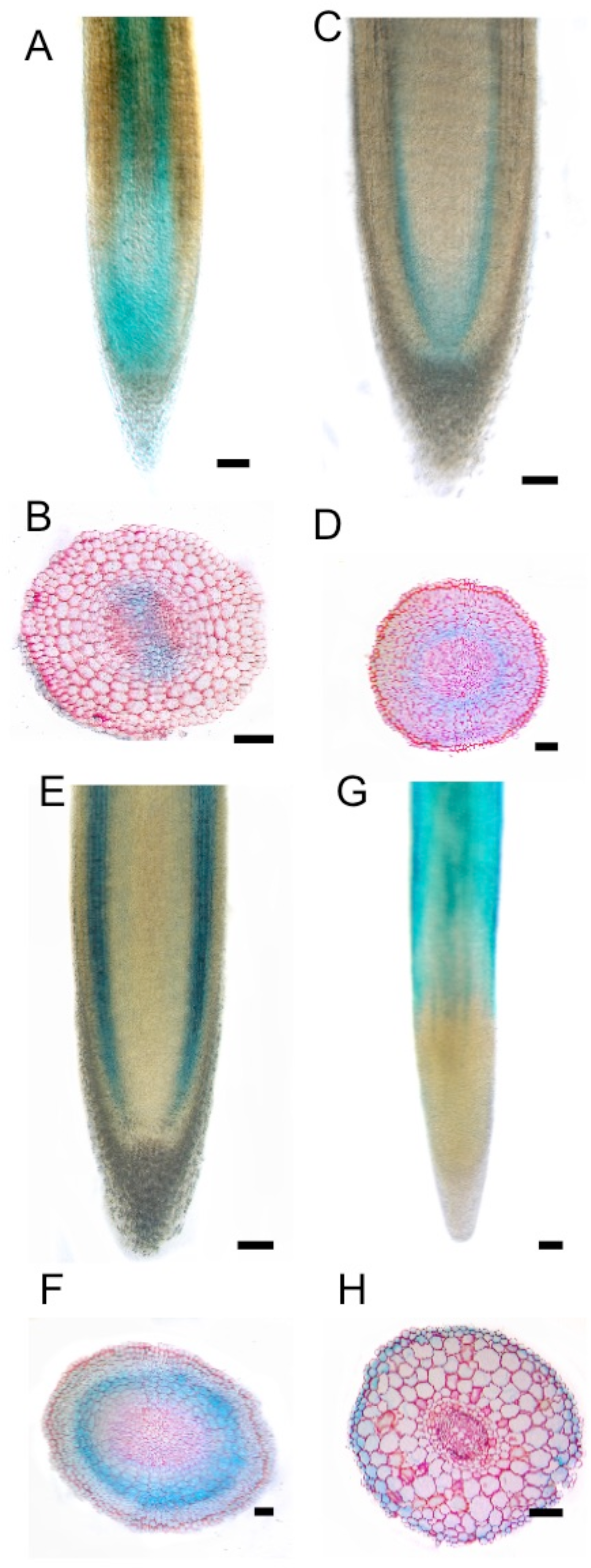
Expression profiles of four white lupin promoters in the second order root. Promoter profiles shown are: *pLaWOL* (A, B), *pLaSCR1* (C, D), *pLaPEP* (E, F), *pLaEXP7* (G, H). Images are longitudinal sections in the meristematic and elongation zones of the tip of the cluster root (A, C, E, G). Cross sections were performed through different cluster root zones: the meristematic zone (D), the elongation zone (F), the differentiation zone (B, H). Scale bars are 100 µm.

### Tissue specific marker expression throughout rootlet primordia development

The various stages of rootlet primordia development were studied from the early divisions up to their emergence in order to determine when tissues differentiate and to provide a developmental atlas of rootlet primordia development. We combined thick longitudinal cross sections (Fig. 6) to provide stronger GUS staining together with thin radial cross sections (Fig. 7) for better accuracy. The *pLaWOL:GUS* marker showed strong signal as early as stage II (Fig. 6A) corresponding to 6 pericycle cells in P1/P2 right after the first periclinal division (Fig. 7A). At later stages, the staining pattern becomes limited to a group of cells at the base of the primordium that might be derived from P1 (Fig. 6B,C and Fig. 7B). From stage VI onward, staining is clearly observed in elongated cells at the centre of the primordium (Fig. 6D and Fig. 7C). This pattern of expression remains similar up to after emergence corresponding to the newly developed vasculature (Fig. 6E and Fig. 7D). The *pLaSCR1:GUS* endodermal marker expression is first seen in 5 endodermal cells at stage II when these cells appear to divide (Fig. 6F and Fig. 7E). At stage III and IV, staining is not only observed in E1 and E2 (Fig. 6G) but also in the overlaying cortical cells (Fig. 7F). At stage VI, *pLaSCR1:GUS* expression starts to be restricted to two inner layers that are surrounding the stele with a typical U-shaped profile (Fig. 6H). A group of cells abutting these two layers also show weak GUS staining at the tip of the rootlet primordium (Fig. 7G). This expression profile remains the same when the rootlet is about to emerge (Fig. 6I,J and Fig. 7H). Marker line *pLaPEP:GUS* is not expressed in the first stages of rootlet development but is present in the parental cluster root cortex (Fig. 6K-M and Fig. 7I). At stage V, a faded expression can be observed in the tip of the primordium on thin sections (Fig. 7J). This faint expression profile persists at stage VI (Fig. 7K). After emergence, staining is expressed in 3 to 4 cortical cell layers at the edges of the primordium but is not expressed in younger cortical cell files reminiscent of the expression pattern in cluster roots (Fig. 6N,O, Fig. 7L).

**Fig. 6.**
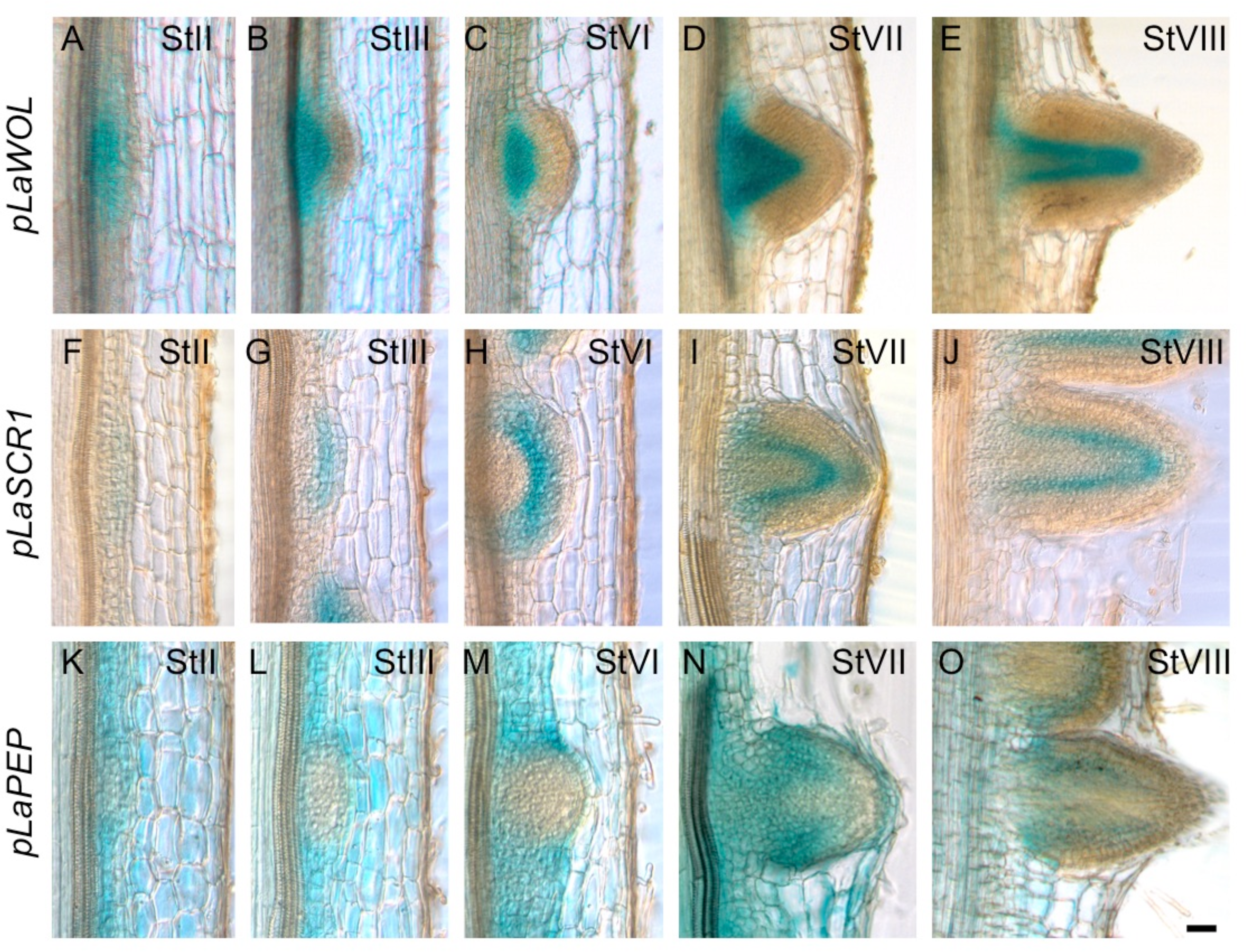
Expression of tissue-specific molecular markers during rootlet primordium development on thick longitudinal sections through cluster roots. Images are 80 µm thick sections showing GUS activity in 3 marker lines. (A-E) *pLaWOL:GUS*. (A) StII. Staining is seen the pericycle. (B, C) StIII, StIV. Staining is observed in the stele-derived tissues. (D,E) StVII, StVIII. Staining is observed in central tissues at the core of the rootlet primordium. (F,J) *pLaSCR1:GUS*. (F,G) StII, StIII. Staining is observed only in the endodermis. (H-J) St VI, VII, VIII. Staining appear to be expressed in layers surrounding the stele. (K-O) *pLaPEP:GUS*. (K,L) StII, StIII. Staining is seen in the cortex of the cluster root but is not expressed in the primordium. (M) StVI. Staining appears in the cortical cells overlaying the primordium. (N,O). StVII, StVIII. Staining is observed in both sides of the primordium but the region at the tip is unstained. Scale bar: 50 µm (A-O).

**Fig. 7.**
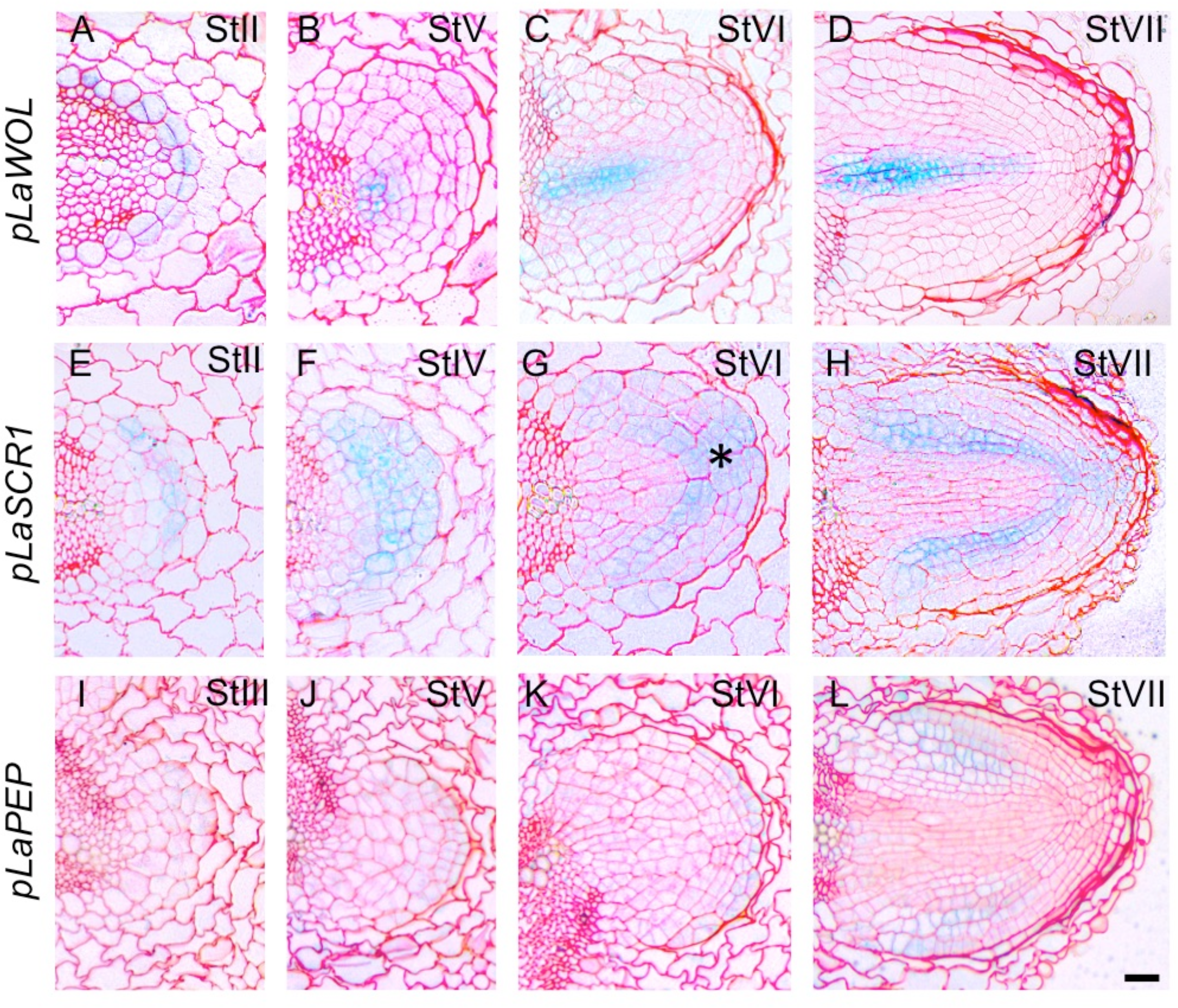
Expression of tissue-specific molecular markers during rootlet primordium development on thin longitudinal sections through cluster roots. Images are 5 µm thin sections showing GUS activity in 3 marker lines. (A-D) *pLaWOL:GUS*. (A) StII. Lupin promoter is specifically expressed in pericycle cell opposite xylem pole, 6 of which seem to be dividing. (B). StV. Staining is apparent in few cells at the base of the primordium. (C, D) StVI, StVIII. Staining is observed in elongated cells found at the core of the primordium. (E-H) *pLaSCR1:GUS*. (E) StII. Staining appear specifically in 6 endodermal cells that appear to be dividing. (F) StIV. GUS staining appear in E1/E2 endodermal layers and cortical cells. (G, H) StVI, StVII. Staining is restricted to two layers around central stele and a group of cells at the tip of the rootlet (asterisk). (I-L) *pLaPEP:GUS*. (I) StIII. GUS staining is difficult to observe at this stage. (J, K) StV, StVI. Weak staining in cortical cells at the tip of rootlet. (L). GUS staining is observed in tissues at the edges of the primordium. Scale bar: 50 µm (A-L).

## Discussion

Previous histological work has described cluster root development in white lupin, showing that rootlets seemingly arise from divisions of pericycle cells facing xylem poles (Johnson *et al*., 1996; Hagström *et al*., 2001; Watt and Evans, 2003). Here, we provide a detailed anatomical study of rootlet development by taking advantage of molecular markers via genetic transformation of white lupin roots (Uhde-Stone *et al*., 2005). Our study reveals that several tissues divide to participate to the formation of the primordium as shown by the cell cycle marker *pAtCYCB1;1*. As in lateral root formation of most species, divisions in the pericycle cells initiate the formation of new rootlets, but several rounds of divisions in endodermal and cortical cells are also observed. These are reminiscent of lateral root formation in other Legumes such as Medicago (Herrbach *et al*., 2014) and other angiosperms (Torres-Martínez *et al*., 2019). It is not entirely clear to which extent these divisions in the outer tissues contribute to the primordium itself or to a protective layer whose fate remains undescribed in white lupin. The presence of more cell layers than in the model plant Arabidopsis, where most of the real time description and cell lineage work has been performed, suggests that different mechanisms may be involved. These mechanisms may not be solely mechanical (stronger pressure, distinct cell wall remodelling processes) but may also include active cell divisions to accommodate the passage of the rootlet primordium. The existence of a temporary cap-like structure has been reported in other species (Torres-Martínez *et al*., 2019) and the numerous divisions in the endodermis seem to generate such structure as can be seen on a Stage VI rootlet primordium (Fig. 1D). Further cell lineage studies would be needed to clearly demonstrate to which extent these dividing cells from outer layers (endodermis and cortex) participate to the primordium itself and hence to the rootlet formation.

Within the rootlet primordium itself, cell division patterns (from the initiation in the pericycle up to the establishment of an organized meristematic zone) is fairly similar to what has been described in other plants and led us to separate its development into 8 stages, similarly to the model plant Arabidopsis (Fig. 8). Although this staging could be seen as somehow artificial due to the numerous extra division events in a multi-layered structure, we believe that it will help comparing similar developmental stages between plant models, especially in the context of transcriptomic analysis similar to the ones already described in Arabidopsis (Voß *et al*., 2015). Interestingly, a meristematic structure is put into place during rootlet development and several divisions occur at the final emergence stage VIII (Fig. 2E) but rootlet development appears to be determinate at later stages (Watt and Evans, 1999). This implies that the primordium meristematic structure, although apparently fully functional at the stages described here, loses its division abilities and fully differentiates later on. Future work should determine at which subsequent stages this switch operates.

**Fig. 8.**
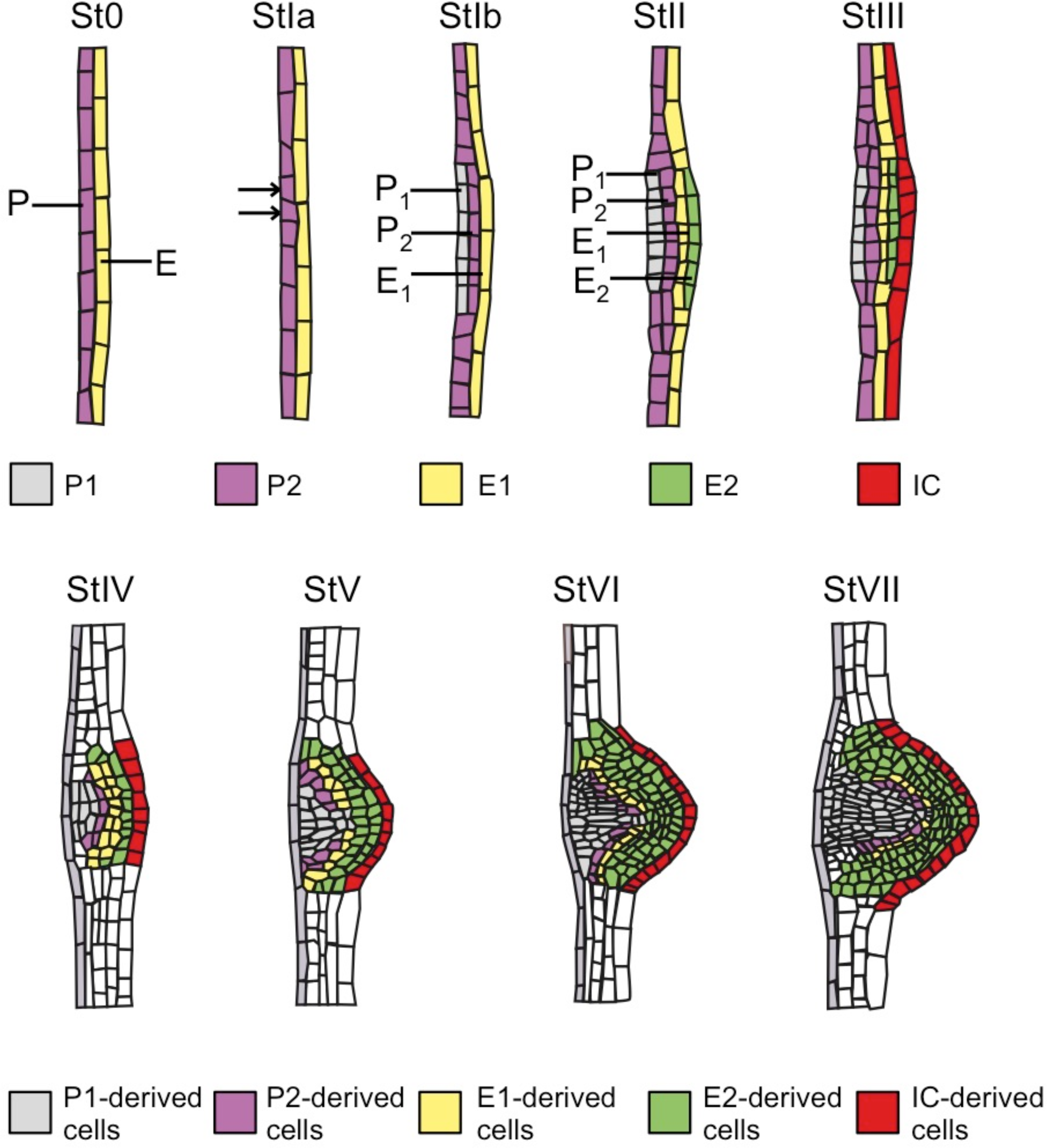
The 8 stages of rootlet development in white lupin. Scheme based on longitudinal sections of white lupin rootlets. Colours indicate putative cell tissues in the growing rootlet primordium from stage I to stage VII. Stage VIII (not represented) would be similar to Stage VII but when the primordium crosses the final layer to reach the rhizosphere. St0: Stage 0, prior to visible divisions, only molecular markers such as *pAtCYCB1;1* can reveal that a site for rootlet formation has been activated. St1a: Stage 1a, first two asymmetrical divisions are seen in pericycle. StIb: Stage 1b, periclinal divisions in pericycle give rise to two layers, P_1_ and P_2_. StII: Stage II, periclinal divisions in endodermis give rise to two layers E_1_ and E_2_. StIII: Stage III, divisions in adjacent cortical cells. StIV: Stage IV, rootlet patterning and tissue differentiation. StV: Stage V, elongation of vascular cells. StVI: Stage VI, differentiation of tissue layers inside rootlet primordium. StVII: Stage VI, further growth of the primordium until it reaches the epidermal layer and prior to emergence (StVIII: Stage VIII).

## Acknowledgements

This project has received funding from the European Research Council (ERC) under the European Union’s Horizon 2020 research and innovation program (Starting Grant LUPINROOTS - grant agreement No 637420 to B.P.). C.G. is the recipient of a fellowship from GAIA doctoral school of Montpellier University. We thank Carine Alcon (PHIV platform) for the help in microscopy and access to imaging facility MRI, member of the national infrastructure France-BioImaging.

## Tables

Table S1. List of gene names and ID numbers, sequence of primers used for cloning.

## Supplementary data

**Fig. S1. Expression profile of the *pAtCYCB1;1:GUS* marker in white lupin cluster root is associated with sites with active cell divisions.** (A) Cluster root tip shows typical spotted expression of dividing cells in the meristematic region. (B) In the differentiation zone, sites of rootlet formation show expression of the marker at early initiation stages. (C) Later on, cell divisions in the rootlet primordium are also seen. (D) Cell divisions can be seen in the active rootlet primordium upon their emergence. Bars are 350µm.

**Fig. S2. Expression profiles of four white lupin promoters in cluster roots.** Promoters profiles shown are: *pLaWOL:GUS* (A-E), *pLaSCR1:GUS* (F-J), *pLaPEP:GUS* (K-O) and *pLaEXP7:GUS* (P-T). Whole-mount view of early differentiation zone (A, F, K and P), late differentiation zone with late stages rootlet primordia (B, G, L and Q), rootlet emergence (C, H, M and R), rootlet midgrowth stage (D, I, N and S) and fully grown rootlets (E, J, O and T). Scale bars are 150 µm (column 1-4), 500 µm (column 5).

